# Consensus rank orderings of molecular fingerprints illustrate the ‘most genuine’ similarities between marketed drugs and small endogenous human metabolites, but highlight exogenous natural products as the most important ‘natural’ drug transporter substrates

**DOI:** 10.1101/110437

**Authors:** Steve O’Hagan, Douglas B. Kell

## Abstract

We compare several molecular fingerprint encodings for marketed, small molecule drugs, and assess how their <u>rank order</u> varies with the fingerprint in terms of the Tanimoto similarity to the most similar endogenous human metabolite as taken from Recon2. For the great majority of drugs, the rank order varies <u>very greatly</u> depending on the encoding used, and also somewhat when the Tanimoto similarity (TS) is replaced by the Tversky similarity. However, for a subset of such drugs, amounting to some 10% of the set and a Tanimoto similarity of ~0.8 or greater, the similarity coefficient is relatively robust to the encoding used. This leads to a metric that, while arbitrary, suggests that a Tanimoto similarity of 0.75-0.8 or greater genuinely does imply a considerable structural similarity of two molecules in the drug-endogenite space. Although comparatively few (<10% of) marketed drugs are, in this sense, <u>robustly</u> similar to an endogenite, there is often at least one encoding with which they <u>are</u> genuinely similar (e.g. TS > 0.75). This is referred to as the Take Your Pick Improved Cheminformatic Analytical Likeness or TYPICAL encoding, and on this basis some 66% of drugs are within a TS of 0.75 to an endogenite.

We next explicitly recognise that natural evolution will have selected for the ability to transport <u>dietary</u> substances, including plant, animal and microbial ‘secondary’ metabolites, that are of benefit to the host. These should also be explored in terms of their closeness to marketed drugs. We thus compared the TS of marketed drugs with the contents of various databases of natural products. When this is done, we find that some 80% of marketed drugs are within a TS of 0.7 to a natural product, even using just the MACCS encoding. For patterned and TYPICAL encodings, 80% and 98% of drugs are within a TS of 0.8 to (an endogenite or) an exogenous natural product. This implies strongly that it is these <u>exogeneous</u> (dietary and medicinal) natural products that are more to be seen as the ‘natural’ substrates of drug transporters (as is recognised, for instance, for the solute carrier SLC22A4 and ergothioneine). This novel analysis casts an entirely different light on the kinds of natural molecules that are to be seen as most like marketed drugs, and hence potential transporter substrates, and further suggests that a renewed exploitation of natural products as drug scaffolds would be amply rewarded.

## Introduction

Given the overwhelming evidence [1–20] that pharmaceutical drugs must and do exploit endogenous transporters that normally transport biological metabolites, and that normally any diffusion of such drugs through the phospholipid bilayer portions of undamaged biological membranes is negligible [1; 3; 5–7; 10; 11; 13; 21], we [2; 22–24] and others (e.g. [16; 25–30]) have been assessing the extent to which marketed (hence successful) xenobiotic drugs are similar in structural terms to endogenous human metabolites (that we sometimes refer to as ‘endogenites’).

Chemical similarity is a slippery concept (see e.g. [31–35] and below) but, leaving aside descriptor-based vectors [36], it is most commonly assessed by encoding the molecules of interest into one or more fingerprints expressed as bitstrings, then comparing the bitstrings, again most commonly in terms of their Jaccard or Tanimoto similarity [37–39]. Our first detailed study [22] noted that the quantitative (Jaccard/Tanimoto) similarity varied markedly with the different (fingerprint-based) encodings used (and we reproduce the essential and Open Access findings in Fig 1A, below), just as does the appearance or otherwise of ‘activity cliffs’ [40–42]. To a certain degree, the shape of the profiles of rank-ordered drugs vs their Tanimoto similarity to the closest endogenous metabolite were smooth curves that differed somewhat. However, this of itself did not tell us - notwithstanding the numerical variation in Tanimoto similarity with each encoding - whether the <u>rank order</u> of individual drugs themselves was more or less well preserved for each encoding. In other words was the drug that was numerically most similar to an endogenite under the MACCS encoding also most similar under (say) the Atom Pair encoding?

**Figure 1.**
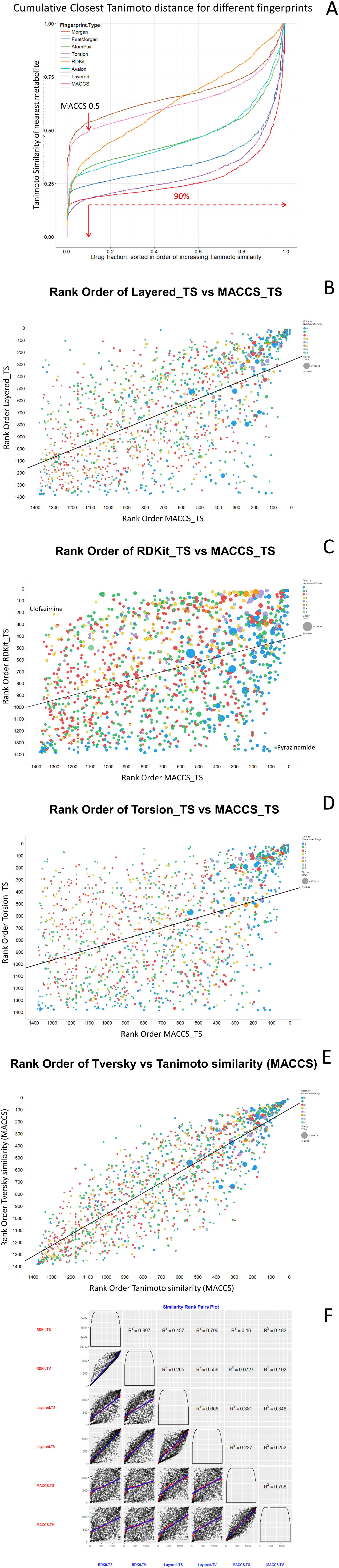
Cumulative similarity and rank order of various encodings. **A**. Cumulative rank order (most similar on the right) of a drug to its closest endogenite for a series of encodings, Redrawn (under a CC-BY license) from [22]. For the other parts of this figure, each symbol represents the rank order of the encodings specified. In addition, although there were no observable trends, symbol size encodes total polar surface area, while colour encodes the number of aromatic rings in the drug (0 blue, 1 emerald, 2 red, 3 yellow, 4 lilac, 5 orange, 6 sapphire, 8 cyan), and these can help to identify individual molecules in different encodings. **B**. Layered vs MACCS encoding, Tanimoto similarities, r^2^ = 0.38. **C**. RDKit vs MACCS encoding, Tanimoto similarities, r^2^ = 0.16. **D**. Torsion vs MACCS encoding, Tanimoto similarities, r^2^ = 0.20. **E**. Tversky (α =0.2, β =0.8) similarity vs Tanimoto similarity for MACCS encoding. r^2^ = 0.76. **F**. Plot of multiple comparisons (blue best linear fit, red best LOESS fit).

The Tanimoto similarity is a true metric, and while it returns a numerical value between 0 and 1 the question also arises as to which values of the Tanimoto similarity genuinely count as ‘significantly similar’ [34] from a utilitarian point of view. Unlike QSAR and other ‘supervised’ methods where there is an objective function, for which the predictions of the model can be tested on unseen data (e.g. where a Tanimoto similarity of 0.85 to a ‘hit’ in a drug discovery assay increases the chance of another hit by 30-fold [43]), the pure notion of chemical similarity is really an ‘unsupervised’ method, and its numerical value is simply that.

Previously, apart from the addition of vitamins, we were rather restrictive about what might constitute an endogenous metabolite or ‘endogenite’, and we here recognise that this restriction was not only unnecessary but potentially very misleading, as any natural molecule with a high k_cat_ or k_cat_/K_m_ [44–47] for a particular transporter might reasonably be regarded as a ‘natural’ substrate for it. In particular, we may suppose that there are or have been natural, bioactive/psychoactive dietary and medicinal components (and their and other microbiome-derived products) that are both beneficial and common enough that the host has essentially been exposed to them more or less regularly through evolutionary time, albeit they do not appear in the common models of <u>human</u> metabolism. Since useful bioactivity in tissues implies uptake, natural selection would then ensure that we had actually evolved transporters for them, and that these molecules, despite not being synthesised by the host, are properly to be seen as ‘natural substrates’ of such transporters. L-ergothioneine is a particularly clear example of this.

### Some mammalian transporters with known selectivity for exogenous natural products

L-ergothioneine (2-mercaptohistidine trimethylbetaine; IUPAC name (2S)-3-(2-Thioxo-2,3-dihydro-1H-imidazol-4-yl)-2-(trimethylammonio)propanoate) is not synthesised by mammals, but exists in a wide range of foodstuffs (especially mushrooms) and may be highly concentrated in mammalian tissues [48; 49]. Several types of evidence imply that it has an important role *in vivo* as a natural antioxidant [50; 51]. First this activity may be measured directly [52–54]. Secondly, decreasing it leads to the accumulation of the products of the interaction of macromolecules with hydroxyl radicals [55–57] and a decreased lifespan in model organisms [58]. Thirdly, it acts as a cytoprotectant [48; 59–63]. Our interest in it here comes from the fact that it has been found to be the natural (or at least most active) substrate of the concentrative, Na^+^-dependent transporter SLC22A4 [64–66] (once referred to as OCTN1, now the ergothioneine transporter, which is also capable of transporting drugs such as the antidiabetic metformin [67]) (for SLC terminology see [68] and http://boparadigms.org/). It was known that OCTN1 transported organic cations, but not what the ‘natural substrate’ might be. Thus, Gmndemann and colleagues [64] incubated cells with and without recombinant OCTN1 transporter expression in paired assays with diluted plasma (taken to contain all candidate substrate molecules) and compared differences in the uptake of the various compounds by mass spectrometry. The first substance identified was proline betaine (stachydrine). Subsequent tests on structurally related molecules showed that ergothioneine was much the best substrate, with an uptake activity almost 100-fold higher than those for tetraethyl ammonium and carnitine [64] (that were previously believed to be the ‘main’ substrates), and that cells lacking the transporter were virtually impermeable to ergothioneine. Since it does not seem to be <u>essential</u> for the growth of the host it has not attained the status of a vitamin, but it is clearly <u>highly beneficial</u>. (Its presence in almost all foodstuffs means that starvation for it specifically, the usual means of discovering or identifying a vitamin, has probably never occurred.) The same may generally be said to be true of other nutritionally beneficial molecules of plant origin, of which the flavonoids are among the best known.

Indeed, in a similar way, it appears that specific transporters for flavonoid-type molecules also exist [69–71], albeit their molecular taxonomy remains unclear [72]. This said, a transporter in plants [73] shows significant homology to bilitranslocase, a liver uptake transporter for blood-derived bilirubin, and bilitranslocase has been shown to transport dietary flavonoids [74], in particular anthocyanins [75–78]. Thus we are led to the view that we should consider as substrates for mammalian transporters not only the known <u>intermediary</u> metabolites, but also a variety of (mainly plant and microbial) <u>dietary molecules that are bioactive and beneficial,</u> even if not essential. This is because organisms will have coevolved with them for millions of years since ‘animals’ began to consume plants [79–82] and to harbour microbes [83; 84]. Even stronger natural selection may be expected since the time that such plants actually began to be utilised in agriculture [85] or prescribed for medical benefit [86; 87], as in Ayurvedic [88–90] and Chinese Herbal Medicine [89; 91; 92] (ca 5-8000y BP). If this is the case, we would expect to find even more structural similarities between drugs and such natural products when these are compared to drug-endogenite similarities (and actually this proved to be the case in a pilot study; Fig 5C of [22]). One purpose of the present paper was to test this idea explicitly and in much more detail. Indeed, it transpires (see a detailed analysis in the body of this paper) that many of the least human-endogenite-like marketed drugs are <u>considerably</u> closer in structure to common plant and microbial secondary products than they are to endogenites. If we take the term ‘natural substrate’ to mean a substance to which an organism has been exposed and for which a transporter has a particularly high k_cat_ or k_cat_/K_m_, it is reasonable to refer to such a molecule as a ‘natural substrate’, as in the ergothioneine/SLC22A4 example above. Another exogenous molecule that seems similarly valuable to mammals [93–104] and other organisms [105], albeit its uptake transporter is not yet known, is pyrroloquinoline quinone (PQQ) [106], also known as methoxatin, a redox cofactor normally associated with prokaryotes [107; 108].

### How similar is similar?

Although the concept of what counts as ‘significantly similar’ must be recognised as highly important in cheminformatics, there has been surprisingly little work done on it; most of it has involved assessing the likelihood that a given similarity could be achieved from a (more or less random) distribution of chemicals [34; 109–113]. Given that our original, underlying interest is in understanding those features of drug and endogenous metabolite structures that tend to determine whether a drug is closer to or far from being most similar to a specific endogenous (or other) metabolite, the question is important (but the distribution of chemical structures is far from random, one having been selected by natural evolution, the other via the processes of drug discovery). Thus, the first part of the present paper analyses that question. The conclusion is that the rank order is reasonably preserved for only a small fraction - some 5-10%-of those drugs that are most similar to an endogenite, but that for the vast majority of drugs not only the numerical value of the (Tanimoto) similarity but also the rank order depends <u>very strongly indeed</u> on the encoding used. However, the fraction of drugs for which different fingerprinting methods of encoding do give consensual answers (Tanimoto similarity ≥ 0.8, for instance) provides a defensible cut-off for what really counts as ‘significantly similar’. This leads to a second part, where we establish that plant-and microbially derived natural products have a much greater similarity to marketed drugs than do the endogenous metabolites of Recon2, and that they are in fact almost certainly the more common ‘natural’ substrates of the transporters on which pharmaceutical drugs hitchhike. This has profound implication for our understanding of the nature and evolution of human drug transporters.

## Experimental

As previously [22–24; 114], we used the list of 1381 marketed drugs and 1113 Recon2-based endogenous metabolites as provided in the Supplementary information to [22]. A number of natural products and other databases exist [115–120]. We have here used the dataset for measured serum metabolites kindly provided by Prof David Wishart and colleagues [121], but removed all substances marked as drugs or that were in recon2. In addition, where noted, we also studied datasets such as UNPD http://pkuxxj.pku.edu.cn/UNPD/ [122] and ZINC [123; 124]. We also obtained a license for the (commercial) Dictionary of Natural Products [125] http://dnp.chemnetbase.com/intro/. All comparisons were done using KNIME-based workflows ([126–128] and www.knime.org/). and in particular we made use of the RDKit nodes [112; 129] (http://rdkit.org/).

## Results and Discussion

### Variance in ‘similarity’ with different fingerprint encodings

Leaving aside molecules that are actually both drugs and metabolites, some drugs are clearly much more similar to one or more endogenous metabolites than are others, and this is true for a variety of fingerprint encodings [22–24] as provided via RDKit [112; 129]. The question thus arose as to whether these similarities extended to the actual <u>rank orders</u> of the drugs (with 1 always being the drug most similar to an endogenite). In other words, was the drug that was most similar to an endogenite when these were represented using the MACCS encoding also most similar with say the Atom Pair encoding? For ease of assessment, Fig 1A recapitulates the original analysis [22] (freely available under a CC-BY license). The three encodings that seemed to maximise the endogenite-likeness of marketed drugs in the earlier paper [22] were the MACCS, Layered and RDKit encodings in RDKit. Thus Fig 1B and 1C show, respectively, the relative rank orders of Layered and RDKit vs MACCS, all using Tanimoto as the metric of similarity. It is clear that while a small subset of the most endogenite-like drugs preserve their rank order between encodings, the rank order for the vast majority depends very strongly on the encoding used (cf. [112]). Also shown for Layered and RDKit (Figures 1B, 1C) are the names of a few drugs for which the differences in rank order are most extreme. The same kind of phenomena are true for Torsion vs MACCS (Fig 1D) and indeed for all the other comparisons tested (data not shown, but all of these data are provided as a spreadsheet via the Supplementary information (DvsMDrugRanks_Full_w_descriptors_hits_with_MACCS_TS.xls)). Overall, while the generation of fingerprints is entirely deterministic, we could discern no real molecular properties that would predict which TS values for a given drug would be ‘high’ or ‘low’ for the set of endogenites. This could be seen as giving weight to view that each is of value and might be used as required.

We also compared the Tversky similarities (α =0.2, β =0.8) (see [24; 114]) for the different encodings, with Fig 1E illustrating its comparison with the rank-ordered Tanimoto similarity for the MACCS encoding. It may again be concluded that while some drugs appear numerically similar to a given metabolite under the different metrics, many do not. However, in this case the correlation (r^2^ = 0.76) is considerably better than that for comparisons of the different encodings. Finally, we illustrate several correlation plots together (Fig 1F).

To encapsulate and to summarise all of the RDKit encodings used in one graph, we compared the <u>sum</u> of all the rank orders with their range (Fig 2A). Thus those at the top right of the plot (Fig 2B) are those drugs that are reliably of high rank order (most similar) whatever the encoding; there are only 44 where the cut-off was (somewhat arbitrarily) drawn. Similarly, a small subset are reliably of low rank order whatever the encoding (and include in particular ‘drugs’ such as fluorinated inhalational anaesthetics that are clearly very far from endogenites) (Fig 2B). Another subset (arbitrarily picked and illustrated in Fig 2C) contains drugs that are mainly not seen as very endogenite-like except in one or two encodings. However, it is obvious that for the vast majority of other drugs the rank order (and hence endogenite-likeness) depends very strongly upon the exact encoding used. For these, endogenite-likeness is not therefore a property of the drug *per se* but additionally (even particularly) of its encoding into whichever fingerprint is chosen. By contrast, the top 10% or so of drugs, that are within a MACCS Tanimoto similarity of ~0.8 to at least one endogenite, are relatively robust to the different encodings (Fig 2D), and one could argue that this relative independence from the nature of the encoding does seem to be a good metric of “similarity”. Although this is something of a self-fulfilling prophecy, inspection of those drugs also clearly <u>does</u> show a metabolite-likeness, especially to endogenites such as nucleobases and sterols (Fig 2E). In a similar vein, although <u>individual</u> encodings can vary significantly, there is a good correlation between the <u>average</u> Tanimoto similarity (for the Morgan, FeatMorgan, AtomPair, Torsion, RDKit, Avalon, Layered, MACCS and Pattern encodings in RDKit) and the drugs that have a Tanimoto similarity ≥ 0.8 in the MACCS encoding (our standard benchmark) (Fig 2F). In Fig 2F, the overall correlation (r^2^) = 0.77 (slope = 0.75), and the variance is much less than that of the rank order.

**Figure 2.**
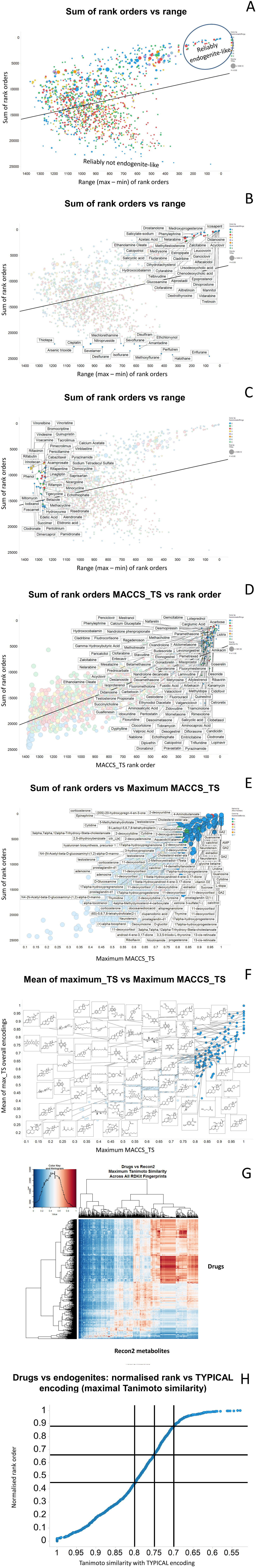
Relationship between the sum and the range of the rank order of the different encodings. A. Overview of the shape of the plot. B. names of marketed drugs that are reliably most or least like an endogenite whatever the encoding. C. Names of drugs for which there is at least one reasonably high rank order but for which mostly they are not encoded as that endogenite-like. D. Names of top 138 drugs (for which TS ≥ 0.8) judged by MACCS similarity in rank order. E. Names of metabolites most similar to the most metabolite-similar 138 drugs (for which TS ≥ 0.8) as judged by MACCS similarity. F. Average value of TS for multiple encodings vs MACCS-encoded Tanimoto similarity. G. Heatmap of similarities of drugs vs endogenites using the TYPICAL encoding. H. Cumulative plot of heatmap data of G.

### Choosing the closest encoding for each comparison

Inspection of figures 2A-2C shows a very considerable range for the majority of molecules, implying that for each molecule there is at least <u>one</u> encoding that is seen as having an especially close value of the Tanimoto similarity for a particular drug-endogenite pair. This best or largest value is here referred to as the Take Your Pick Improved Cheminformatic Analytical Likeness or TYPICAL encoding/similarity. Fig 2G shows a heatmap of the similarities of drugs and metabolites using the TYPICAL encoding (four molecules are dropped because of a curiosity with the Torsion encoding). Under these circumstances, the percentages of drugs having a TYPICAL similarity to an endogenite of 0.8, 0.75 and 0.7 are, respectively, 45%, 66% and 88%, as may also be observed in the cumulative plot of Fig 2G shown in Fig 2H.

### Use of the maximum common substructure

Another means of comparing structural similarities (and hence rank orders), and one that does not depend nearly as much (but see [130]) on the fingerprint encoding used, is according to the size of their maximum common substructure (MCS). As before [24], we have here done this using a series of values of the Tversky similarity, varying the Tversky similarity parameters (α and β) such that their sum was either 1 (Fig 3A) or 2 (Fig 3B). Since the encoding is the same, the correlations between the rank orders for different values of α and β are much higher than for the different encodings, with a clear trend of similarities being visible in the violin plot of Figure 3C. Finally, here, we illustrate a comparison of the MCS with a Dice coefficient (α = β = 0.5) and the MACCS_Tanimoto; again for the drugs with the highest values of TS to a metabolite (we illustrate those over 0.85 this time) there is a clear consistency of the metabolite-likeness of their fingerprint-based MACCS encoding and their MCS with a Tversky similarity.

**Figure 3.**
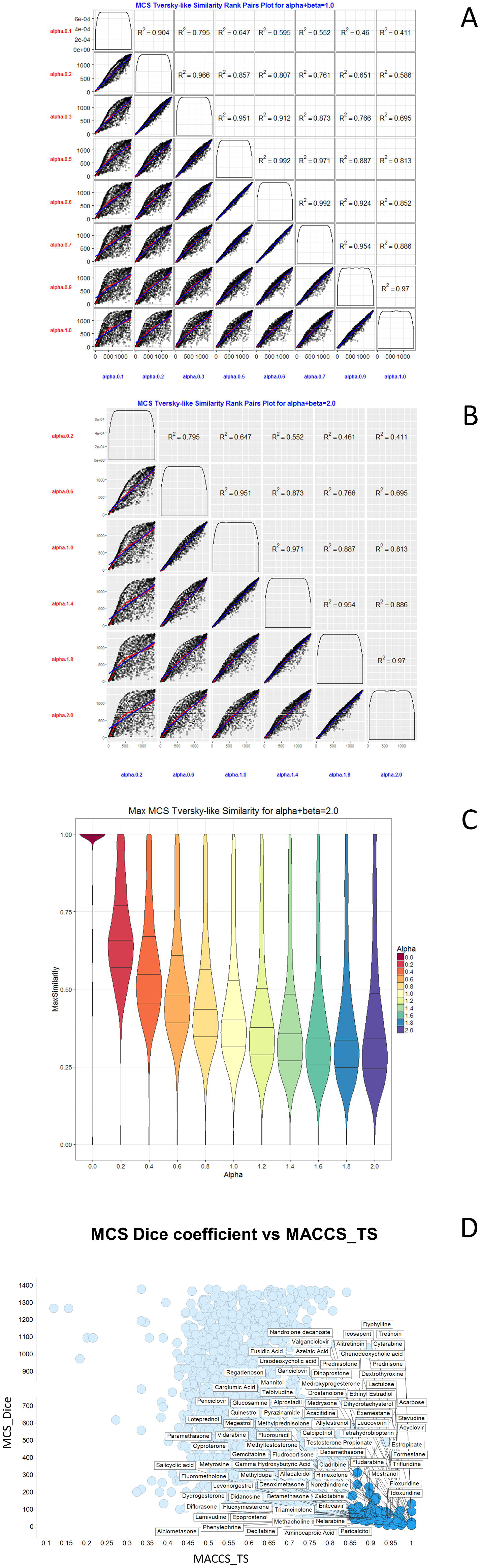
Rank order of drug-endogenite similarities as judged by the size of their maximum common substructures (MCS), for varying values of the Tversky similarities (α and β). A. Sum of α and β = 1. B.Sum of α and β = 2. C. distribution of values of the MCS for varying values of Sum of α and β when their sum is 2. D. MCS Dice coefficient against MACCS Tanimoto similarity.

### Assessment of contribution of descriptors to rank orders using random forest regression

In order to understand the structural bases for some of the rank orders, we set up a random forest regression (see e.g. [131; 132]) to assess whether we can indeed predict the rank of a particular drug molecule in terms of the Tanimoto similarity of its closest endogenous metabolite. As this is a supervised method, we trained on a subset of examples to see if we can predict an out-of-the-box set. The results are shown in Fig 4A, indicating a reasonable degree of success. To ensure that this was not due to any kind of overtraining, we performed target permutation i.e. we randomised statistically the values of the target column one thousand times (data not shown). This served to break any true correlations between the features and the targets, showing that the observed correlations were indeed real. Fig 4B shows an equivalent permutation on the features (i.e. the RDKit descriptors), to assess those which most contributed to the observed correlations. Finally, Fig 4C shows the improvement in correlation that was observed (in out-of-the-box data) as the number of features was increased; evidently ten features were sufficient to achieve the maximum correlation observed.

**Figure 4.**
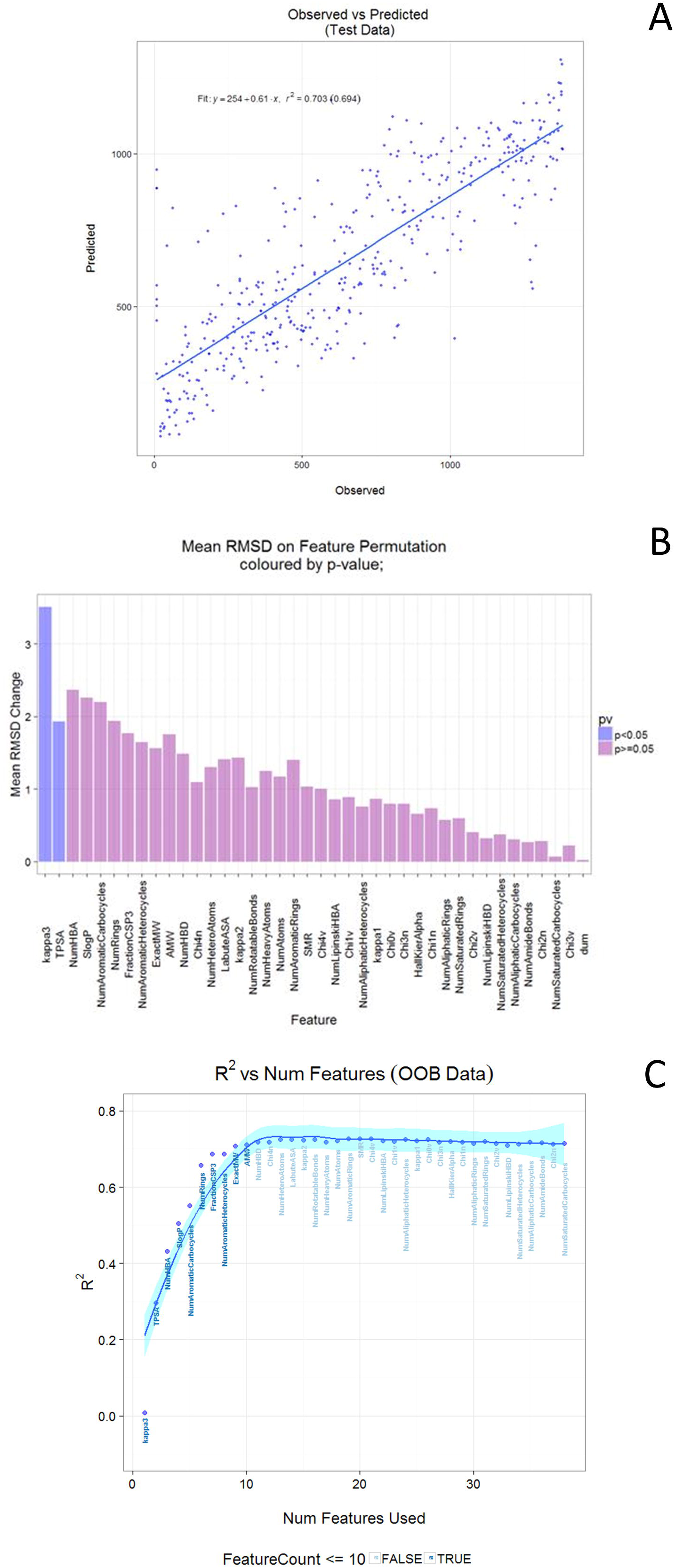
Random forest regression of RDKit features on the rank order prediction of the Tanimoto similarity of a given drug molecule, using the RDKit fingerprint encoding (a graph-based method that should not of itself be related to physicochemical descriptors) and the Tanimoto similarity. A. Predicted against actual. B. Ranked order of relevant features. C. Stepwise improvement in regression as features are added in the order of those seen in B.

### How similar are drugs to dietary and medicinal natural products from plants and microbes?

As mentioned in the introduction, and leaving aside molecules that are actually both drugs and metabolites, some drugs are clearly much more similar to one or more endogenous metabolites than are others, and this is true for a variety of fingerprint encodings [22]. One question that we have not previously asked is about how much better our ‘similarities’ might be if we also used dietary or bioactive molecules that are <u>not</u> in Recon2. Specifically, including for evolutionary reasons rehearsed in the introduction, the question thus arose as to whether these similarities could be increased, especially for the “less similar” drugs, when we began to include bacterial, plant, and fungus-derived secondary metabolites.

We recognise that we must, so far as is reasonable, compare like with like, and certainly it can always be claimed that there is a greater likelihood of finding a molecule with a greater similarity (in a given encoding) as the size of a database is increased *per se.* The normal way of dealing with this is simply to quantify the likelihood that a given similarity could be achieved from a (more or less random) distribution of chemicals taken from that database [34; 109–113]. This is not entirely logical from a biological point of view, however, since such samples are <u>not</u> (from) a random distribution but <u>are the products of evolutionary selection</u>. Thus we prefer other arguments, based on comparing biologically relevant databases. We do, however, also recognise that the gross distribution of properties such as MW, logP, total polar surface area (TPSA) and so on differs between the different databases, and we will need to ensure that this is not a trivial cause of any differences observed. Thus in some cases we used the MatchIt algorithm [133; 134] (and its attendant R code) to select subsets from the various databases with the same distributions of properties as those of endogenites.

The “Universal Natural Products Database” (UNPD) (http://pkuxxj.pku.edu.cn/UNPD) is said [122] to be the largest noncommercial and freely available database for natural products. At the time of its original publication [122] UNPD comprised 197,201 natural products from plants, animals and microorganisms. Our first task here involved regularising or ‘cleaning’ the UNPD for our purposes. Cleaning was perfomed using a KNIME workflow and lowered the number of molecules included from the ca 229,000 initially logged when we downloaded it in December 2016 to 155,048. The main ‘loss’ was due to the loss of (what we could not deconvolve as) duplicates. Some of these may have been stereoisomers, but the 2D connection table provided contained no stereochemistry. Figure 5 shows the distributions of four properties between the endogenous metabolites of Recon2 and the contents [122] of the ‘cleaned’ UNPD natural products database. Although there are clear differences, they are in fact surprisingly similar (see also [22; 122]), and as noted above, individual descriptors had only a minor influence on the random forest model.

**Figure 5.**
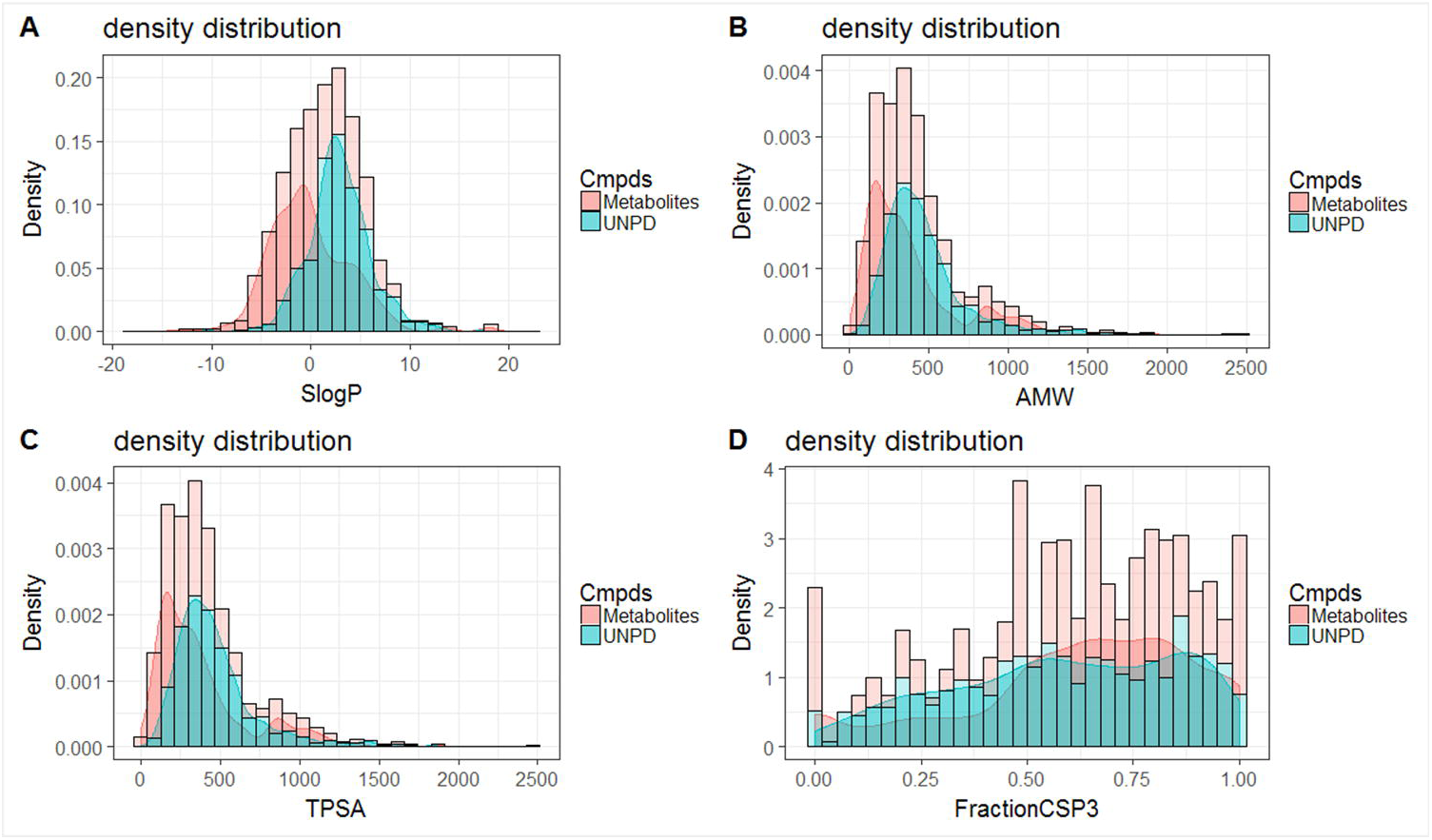
Distribution of four properties between the endogenous metabolites of Recon 2 [135] and the cleaned version of UNPD [122]. The original UNPD file as downloaded contained 229,358 molecules. ‘Cleaning’ removed duplicates as well as molecules that were in either Recon2 or in the list of marketed drugs, both of which were precisely as described and used previously [22–24]. The resulting spreadsheet retained resulted in a total of 155,048 molecules. The smoothed version is the probability density as derived from the R-encoded kernel density estimator at https://www.rdocumentation.ora/packaaes/stats/versions/3.3.2/topics/densitv.

We next (Figure 6A) compared the ordered results of the Tanimoto similarity of the various marketed drugs to those of the nearest representative in our ‘cleaned’ version of UNPD. The results are absolutely striking; while 90% of marketed drugs had an endogenite with a TS > 0.5, the corresponding value for UNPD of 90% was a TS of 0.7. Table 1 shows the %age of drugs with a closest molecule with a TS exceeding various values (MACCS encoding) for endogenites and UNPD library members. Fairly obviously, the chance of finding a close homologue is massively greater (often four-fold or more) for the latter, especially for TS values greater than about 0.7.

**Figure 6.**
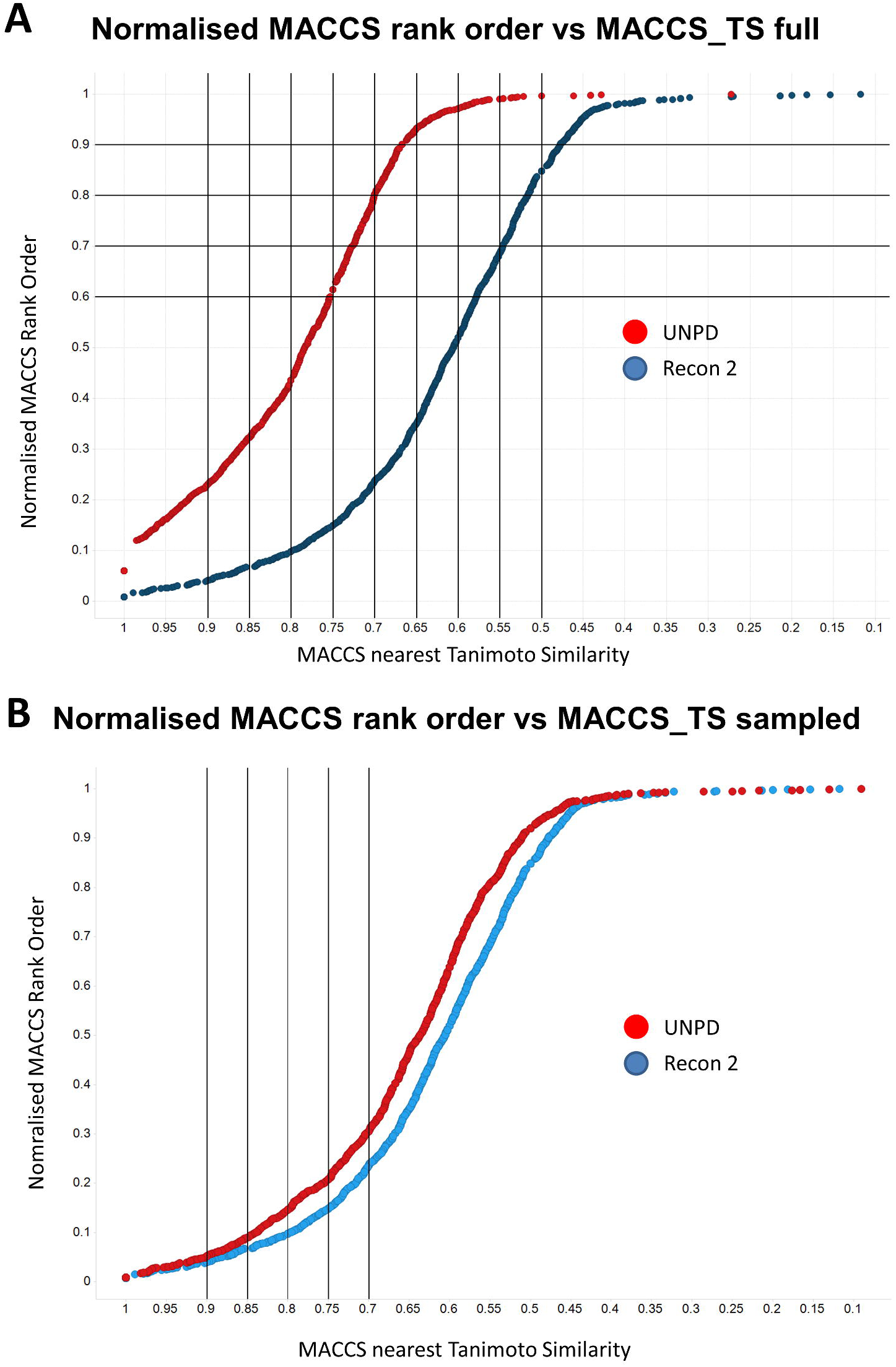
Relationship between the normalised rank order of the nearest database molecule to marketed drugs for (red) the cleaned version of UNPD [122] and (blue) Recon2. ‘Cleaning’ removed duplicates as well as molecules that were in either Recon2 or in the list of marketed drugs, both of which were precisely as described and used previously [22–24]. **A**. Full, cleaned version of UNPD. **B**. A sampled version of UNPD using the same number of molecules as those in Recon2 sampled as per the distributions in Fig 5.

**Table 1.** Tabulation of data from Fig 6A.

| TS > at least | Drugs/endogenites (%drugs) | Drugs/UNPD (% drugs) |
| --- | --- | --- |
| 0.5 | 1185 (85.8%) | 1375 (99.6%) |
| 0.55 | 941 (68.1%) | 1368 (99.1%) |
| 0.6 | 708 (51.3%) | 1339 (97.0%) |
| 0.65 | 486 (35.2%) | 1289 (93.3%) |
| 0.7 | 322 (23.3%) | 1113 (80.6%) |
| 0.75 | 201 (14.6%) | 830 (60.1%) |
| 0.8 | 138 (10.0%) | 614 (44.5%) |
| 0.85 | 93 (6.7%) | 447 (32.4%) |
| 0.9 | 53 (3.8%) | 314 (22.7%) |

One obvious point is that the number of molecules in the cleaned UNPD is roughly 100x greater than the number of those in Recon2, so it could be argued that this alone means statistically that there is simply a greater likelihood of finding a ‘closer’ molecule. While true, this ignores the biology (and the fact is that we did find massively more structurally close natural products than endogenites for a given drug), but we report both analyses. Thus, Fig 6B shows the same comparison as that of Fig 6A save that the UNPD molecules are sampled so as to be numerically equal to those of Recon2, and to share its distribution of the four molecular properties shown in Fig 1. In this case, the ‘advantage’ of UNPD is clearly diminished, albeit still substantial, with 70, 124, 197, 282 and 417 molecules with a TS > 0.9, 0.85, 0.8, 0.75 and 0.7 for UNPD, but only 57, 93, 130, 209 and 329 equivalently for Recon2. Thus for some values of TS, UNPD can enjoy a 50% advantage over Recon2 even when comparisons are strictly scaled to numbers, whatever the biology. We also ran the sampled version multiple times, to look at the ‘range’, but the numbers involved were great enough that this made negligible difference. Note that Recon2 does <u>not</u> contain the thousands of permutations of triglycerides and the like [136], that would increase its size substantially but not provide significantly better hits (i.e. the ‘100x’ figure above is rather a substantial overestimate of the differences in true size), and we also know of many more endogenites that are not yet in Recon2 (see e.g. [9]).

Fig 7A shows a similar comparison for the natural products in a cleaned-up version (see Materials and Methods) of the Dictionary of Natural Products (DNP) [125], seen as a fair comparison [137], and again using the MACCS encoding. Our cleaned DNP (with marketed drugs, endogenites and duplicates removed) contains 72,442 molecules, including 32,390 that are already in UNPD (implying 41,228 that are ‘new’, but also implying 123,443 that are in our UNPD but not in our DNP). Here there are at least 37% of drugs that have a TS greater than 0.8 to the nearest database member, and 50% have a TS greater than 0.75, contrasting with values for Recon2 of just 10% and 15%, respectively. These findings are roughly similar to (but the similarities slightly lower than) those from UNPD, indicating that at least some ‘winners’ are unique to UNPD and some to DNP. In a similar vein, Figure 7B shows the sampled version, with little impact.

**Figure 7.**
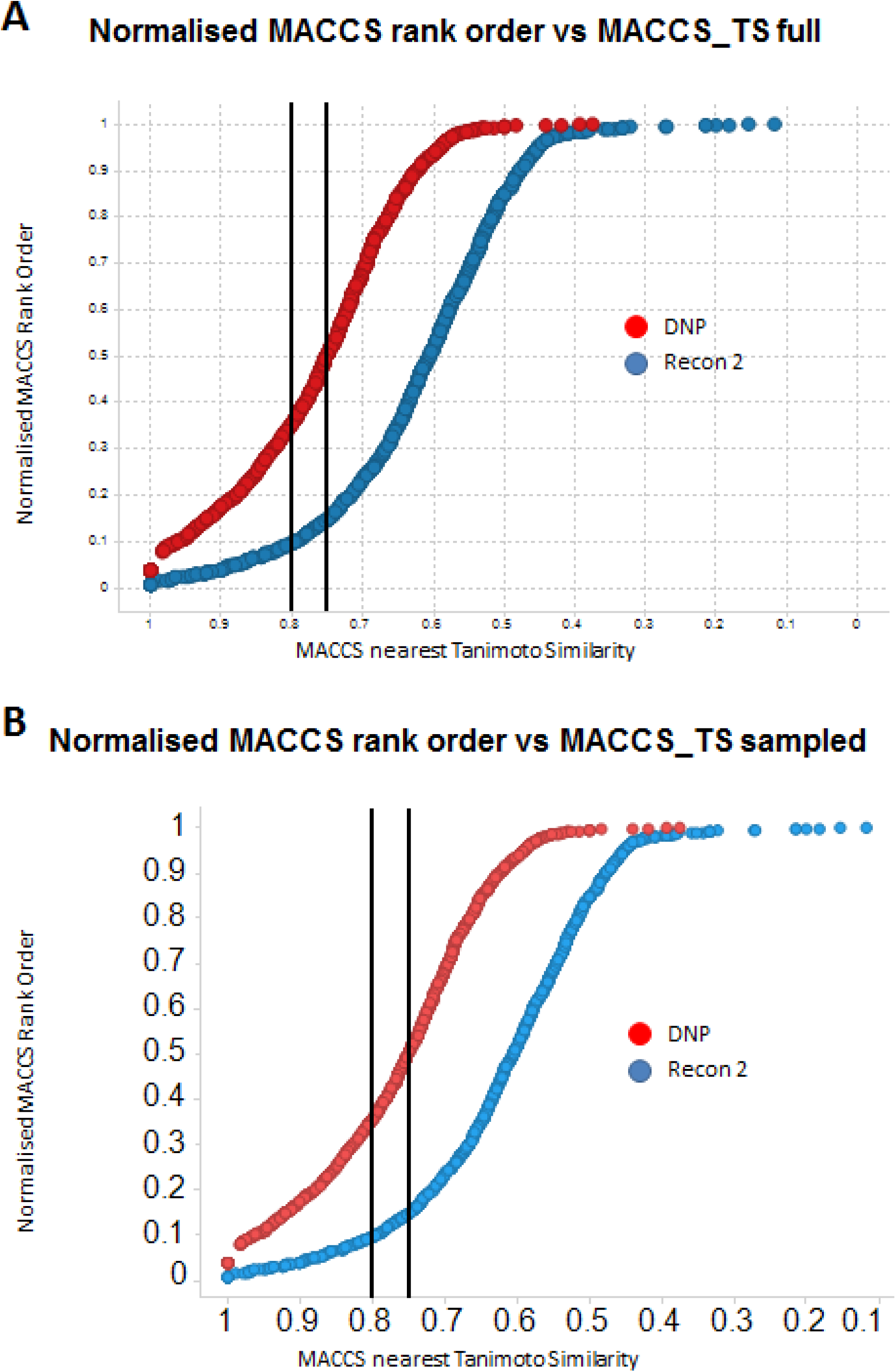
Relationship between the normalised rank order of the nearest database molecule to marketed drugs for (red) the cleaned version of the Dictionary of Natural Products (DNP) [125] and (blue) Recon2. ‘Cleaning’ removed duplicates as well as molecules that were in either Recon2 or in the list of marketed drugs, both of which were precisely as described and used previously [22–24]. **A**. Full, cleaned version of DNP. **B**. A sampled version of DNP using the same number of molecules as those in Recon2 sampled as per the distribution of the properties in Fig 5.

Fig 8 shows the effects of cleaning and the degree of overlap of molecules in our cleaned versions of UNPD and DNP.

**Figure 8.**
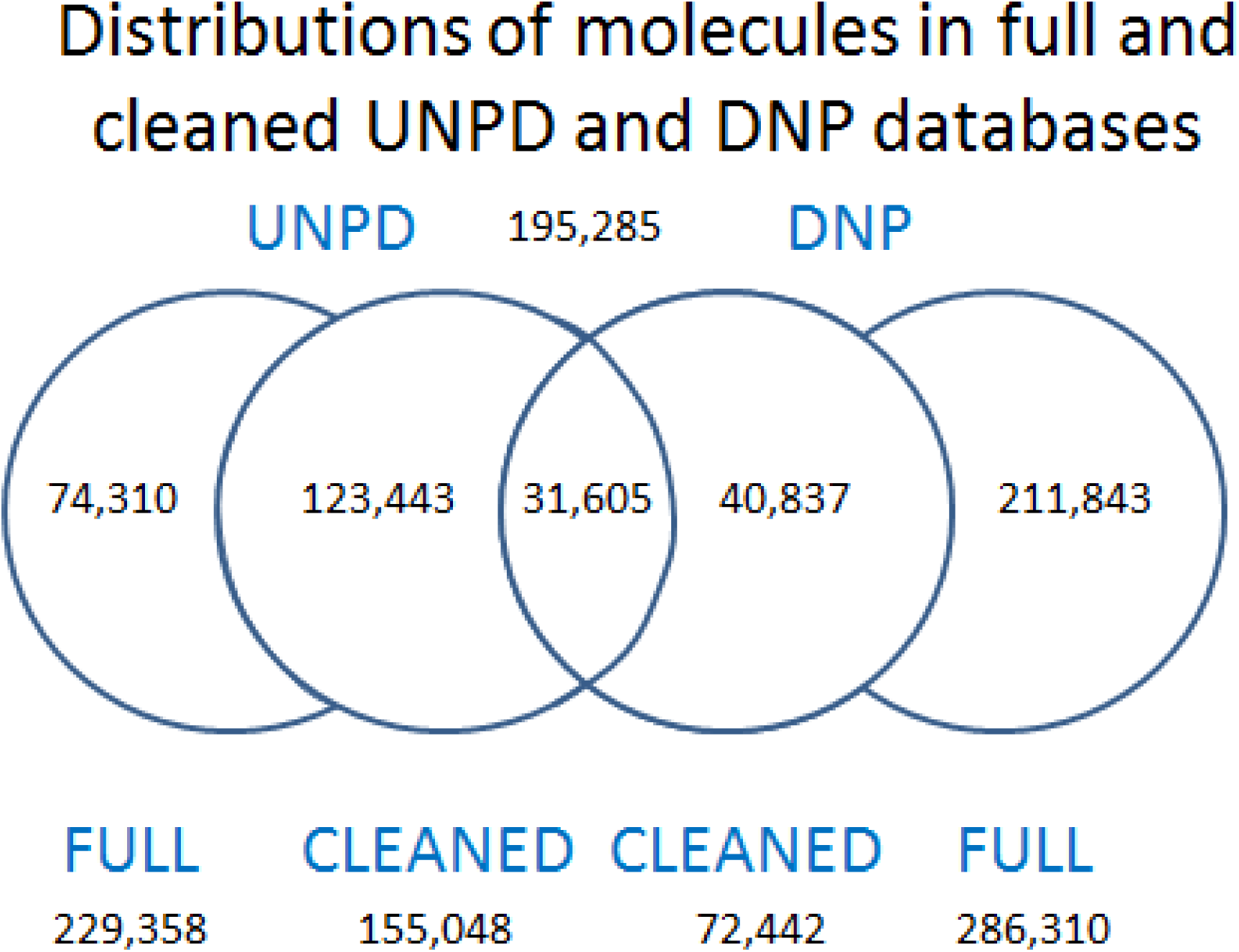
Overlaps between UNPD and DNP databases. ‘Cleaning’ removed duplicates as well as molecules that were in either Recon2 or in the list of marketed drugs, both of which were precisely as described and used previously [22–24].

**Figure 9.**
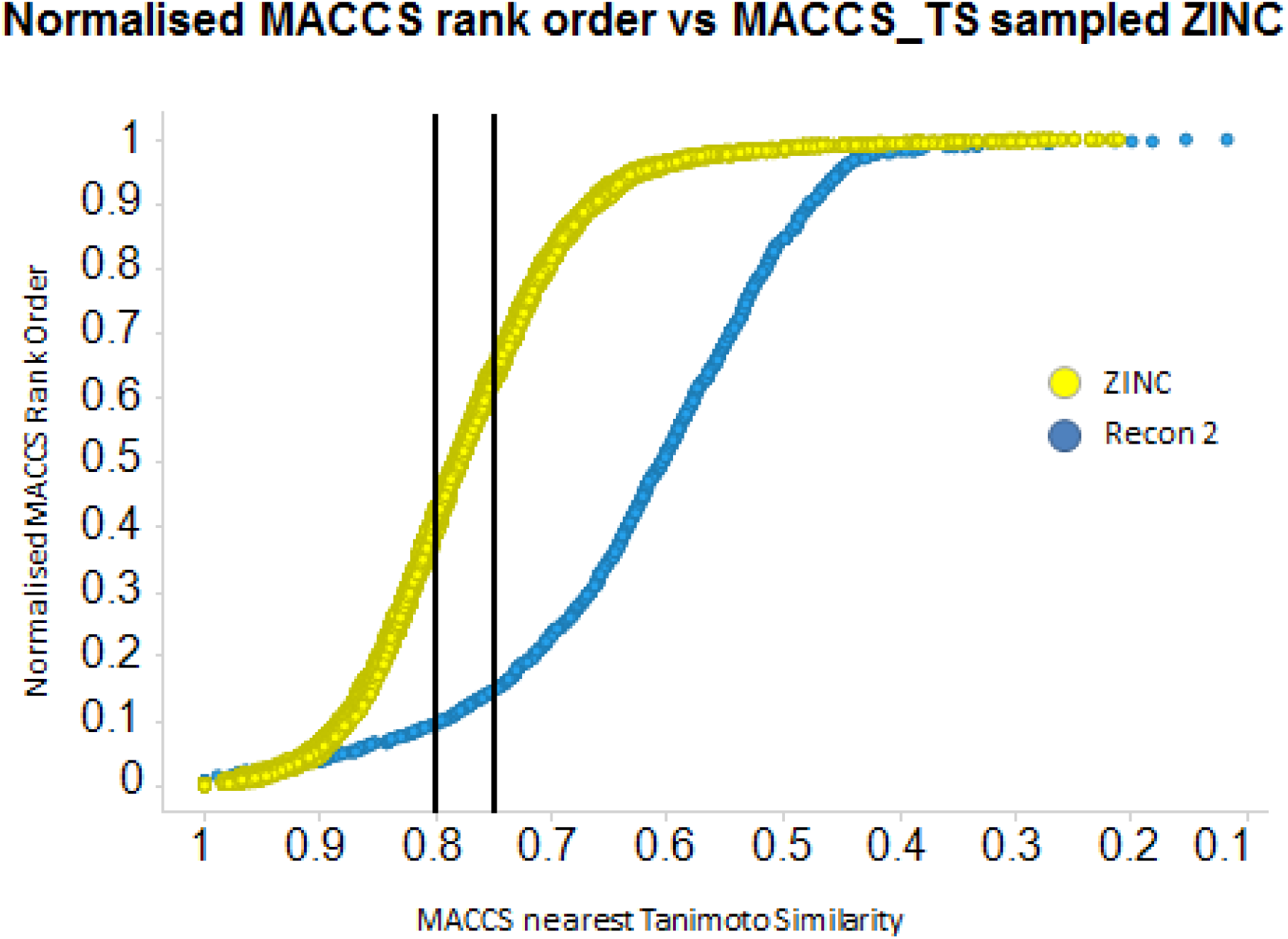
ZINC database. Relationship between the normalised rank order of the nearest database molecule to marketed drugs for the ZINC database (ZINC) [123] (red) and Recon 2 (blue). The ZINC database was ‘cleaned’ to remove molecules that were in either Recon2 or in the list of marketed drugs.

The ZINC database [123] includes a very large number (ca 16M) and variety of synthetic molecules. As usual, we cleaned it to remove any molecules that were marketed drugs or Recon 2 metabolites, and ran it (and recon 2) against drugs as above. This time we ran it as 50 subsets, each of some 148,000, to show the range of curves that we could get. Thus the percentage of ZINC samples that had a member that was within a TS of 0.8 to drugs is between 35 and 45% depending on the sample, while that for a TS of 0.75 or greater varied from 0.59 to 0.74. This implies a considerably greater variation than that for the natural products.

Another source of candidate transporter substrates was the list of molecules observed in serum as catalogued at http://www.serummetabolome.ca/. on the grounds that if they had reached the bloodstream they must have been transported there. We produced a version of this that again lacked all marketed drugs and recon2 metabolites, amounting to some 1480 molecules. Inspection of these indicated that they were mainly nutrients and their metabolites, along with the metabolites of various medicines. Of course what is in serum largely reflects what was recently ingested, and so it can hardly be expected to include all the natural products listed in UNPD and DNP. The curves are shown for both the subset normalized to the size of recon 2 (Fig 10A) and the full set (Fig 1B). Clearly, again, there are a significant number of ‘serum’ molecules that are not in Recon2 yet are structurally closer to drugs. A detailed analysis beyond this is not particularly pointful, since clearly what is in serum reflects recent ingestion only, and this is only a small subset of the contents of UNPD and DNP (Fig 10C).

**Figure 10.**
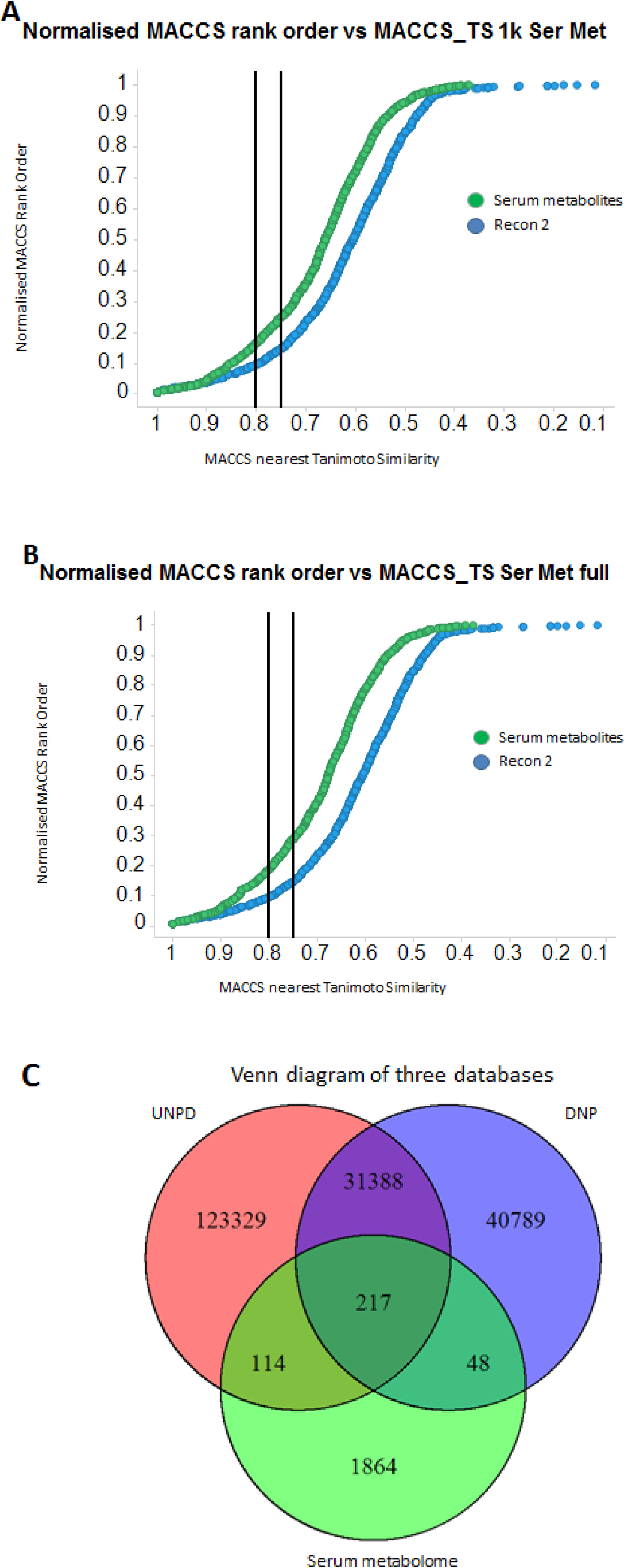
Human serum metabolome. Relationship between the normalised rank order of the nearest molecule to marketed drugs for the ‘human serum metabolome’ database [121] (green) and Recon 2 (blue). The human serum metabolome database was ‘cleaned’ to remove molecules that were in either Recon2 or in the list of marketed drugs. **A.** Sampled subset to be the same size and property distribution as that of recon2. **B.** Full set of ‘human serum metabolome’ molecules after cleaning. **C.** Venn diagram of the co-distributions of molecules in the cleaned versions of UNPD, DNP and the human serum metabolome databases.

### The union of the UNPD and DNP databases

Since there was (surprisingly) little overlap in the contents of the ‘cleaned’ versions of UNPD and DNP (Figures 8, 10C), it was of especial interest to run the analysis on their union, a set of some 195, 285 molecules. The results are shown in Figure 11A for the full set for the MACCS rank order and Tanimoto similarities, and for each of the standard RDKit encodings in Fig 11B for a 148k subset. The results (Fig 11A, B) are absolutely striking: for the MACCS encoding, 45%, 60% and 80% of marketed drugs are within a TS of 0.8, 0.75 and 0.7 to at least one inhabitant of the union of the UNPD and DNP databases, regardless of the inclusion of recon2 metabolites. Fig 11C shows the data for the multiple encodings. On this basis, 80% of all drugs are within a TS of 0.8 of a natural product for the Patterned encoding, and almost all the ‘missing’ molecules with similarities above say 0.75 are natural products. This becomes even clearer in Fig 11D, where we chose the TYPICAL encoding, i.e. that which maximises the TS between a drug and a comparator molecule, regardless of the encoding, and performed this on the full combined natural proucts dataset of ~196,000 molecules. The result was that 92%, 98% and 99.5% are within a TS of a natural product of 0.9, 0.85 and 0.8, respectively. Fig 11E shows the rather widespread distribution of ‘winning’ similarities between the different encodings. Each is represented at least once, and, interestingly (as is also clear from Fig 11C), ‘patterned’ is the most common. This is a more recent addition to the RDKit stable, and was not available when the comparison in Fig 1 was done [22]. However, the next most used are RDKit, MACCS, Layered, and Morgan; with the exception of the latter, the same may also be inferred from the endogenite-only data in Fig 1. There were exactly 500 occasions on which at least one endogenite was the closest in at least one encoding. However, when the TYPICAL encoding used, <u>each</u> of the 1381 drugs was closer to (or equal with) at least one exogenous natural product that is in either or both of UNPD and DNP than it is to an endogenite (data not shown). Finally, in previous work [15], we had compared a metaanalysis of 680 Caco-2 cell permeabilities of 187 marketed drugs with their endogenite-likeness (finding none). Neither did we find any relationship between Caco-2 permeability and any analysis of closeness to the union of the UNPD and DNP databases; as an illustration, Fig 11F compares the same permeabilities with the maximum pattern TS. Clearly any causal relationship that may exist is overwhelmed by the unknown variance [14] in k_cat_, promiscuity, and transporter expression levels.

**Figure 11.**
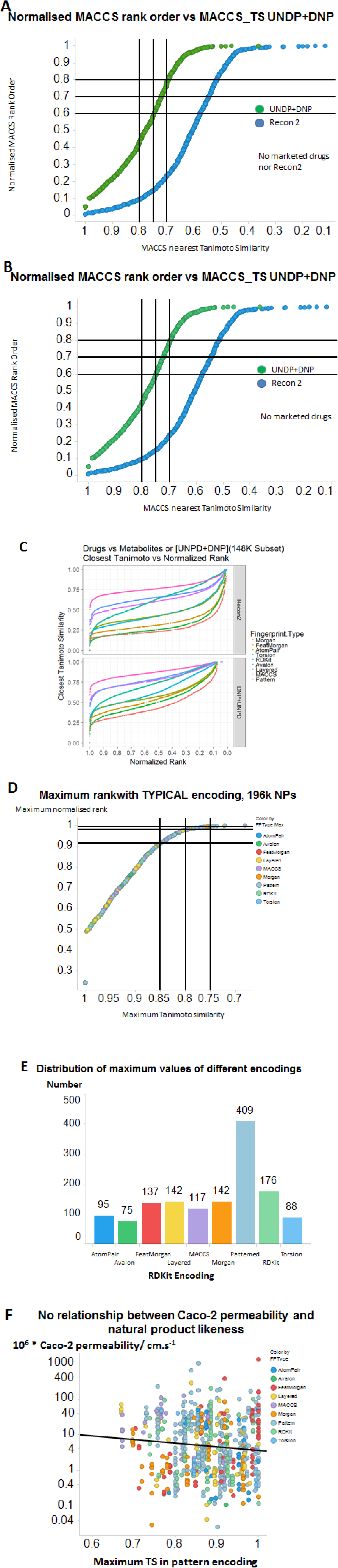
**A**. Relationship between the normalised rank order of the nearest molecule to marketed drugs for the union of the UNPD and DNP natural products databases (green) and Recon 2 (blue). The UNPD and DNP databases were ‘cleaned’ to remove molecules that were in either Recon2 or in the list of marketed drugs. **B.** Similar data plotted for a 148k normalised subset, also lacking marketed drugs. **C**. The same, for each of the standard RDKit encodings. **D**. The same for the TYPICAL encoding. **E**. Distribution of encodings used in the winning molecules that contributed to the TYPICAL encoding. **F**. Lack of relationship between Caco-2 permeability and maximum TS of the union of the UNPD and DNP databases using the pattern encoding (slope =-1.02, r^2^ = 0.011. Drugs are coloured according to the encoding with the largest TS.

## Discussion

### Different similarities from different encodings

As we continue to analyse the structural ‘similarities’ of drugs and endogenites in different ways [14; 15; 22–24; 114], it is becoming increasingly clear that a given drug-endogenite pair can have a highly variable numerical similarity depending on which fingerprint encoding or metric of similarity is used. In the present work, we extend this recognition to the fact that - apart from a very small subset of ‘reliably’ endogenite-similar drugs - the degree of similarity and <u>its rank order</u> can be dominated by the encoding used. This was largely not the case in the analysis of Riniker and Landrum [112], who compared the similarity of fingerprints of larger and very different datasets of library compounds. We also noted that we could predict the rank order using random forest regression, so it was, as expected, a deterministic property.

Willett and colleagues have suggested that ‘fusing’ the results of different fingerprint encodings may give more robust analyses [138–144]. Our strategy is somewhat similar in that we recognise the highly variable rank orders (and Tanimoto similarities) that result from the different encodings, such that their variance tends to increase with their mean rank order. Summing (equivalently, averaging) the rank orders (see also [138; 144]) was a particularly convenient means of combining the data. When this was done, there was a clear trend to the effect that there was much less variance among those molecules with the most reliably high rank order (numerically small values), leading to a conclusion that for Tanimoto similarity values over ~0.75 or 0.8 the similarities are fairly robust to the specific encoding used, and <u>on that basis</u> may reasonably be considered ‘reliable’ or ‘significant’. This said, there was often at least <u>one</u> encoding for which the TS between a given drug and at least one endogenite exceeded 0.75, such that taking the maximum of these <u>regardless</u> of the encoding did increase the number of ‘similar’ endogenites. Unfortunately, with occasional exceptions [145; 146], our knowledge of the substrate specificities of individual transporters is inadequate to the task of assessing whether the ‘nearest’ (or a nearby) metabolite is actually the ‘natural’ or endogenous substrate [12]; by and large, that will have to await further experimentation [12].

### Natural products that are nutrients or bioactive drugs must necessarily be transported

As was recognized from its inception [147], the ‘rule of 5’ [147–153] is taken not to apply to large, natural products, and also does not apply if transporters are involved in the uptake. Natural products have been and remain a major source of successful (marketed) pharmaceutical drugs [154–161], Indeed, about one half of new drugs are based closely on natural products [156; 157], and many transporters exist for them [162–165].

It is to be assumed that anything that is of eventual benefit to (the reproductive fitness of) an organism is likely to be a subject of natural selection and adaptive evolution, even in the laboratory [166], *in vitro* [167], and *in silico* [168]. Thus, if the eating by a mammal of say a plant or fungus has beneficial properties in terms of improving the mammal’s reproductive longevity, selection will act to enhance the uptake of the bioactive principles, at least to a non-toxic level. Certainly, as mentioned, it is well established that natural products themselves contribute importantly to the development of successful drugs (e.g. [154–156; 158–161; 169–172]). Consequently it is clear that a or the ‘natural’ substrate of at least some transporters is in fact likely to be an <u>exogenous</u> molecule that imparts health benefits, and ergothioneine and its uptake by SLC22A4 seem to provide a very clear example [49; 50; 58; 64; 65].

The acquisition of the ability to maintain lactase into adulthood (hence to tolerate lactose and dairy products) is highly heterogeneous and of recent evolutionary origin (~5000y BP [173–177]). Similarly, the actual human selection and prescription of plants as medications is of similar vintage [88; 90–92], albeit hominids and their evolutionary predecessors have been eating plants and fungi for many more millennia (angiosperms appear 50-100My ago). Hence, while it is not possible to replay the evolutionary tape, it is entirely reasonable that many of the several hundred human uptake transporters [68] were in fact selected, at least in part, precisely to transport exogenous secondary metabolites. Indeed, mammalian transporters are well known for their ability to transport many exogenous natural product drugs, e.g., SLCO family members for penicillins [178; 179], cephalosporins [180; 181], tetracycline [182], caffeine, theobromine and theophylline [183] and digoxin [184], SLC22 for berberine [185] and protoberberines [186], morphine [187], erythromycin [188] and theophylline [188], SLC15 for penicillins and cephalosporins [189; 190], SLC6 family (norepinephrine transporters) [191] for ephedrine derivatives [192], and SLC36 for arecaidine (an active constituent of the *Areca* nut, often wrongly referred to as the betel nut) [193].

Of necessity, there are transporters for exogenous natural products that serve as vitamins, such as ascorbate (SLC23 family [194; 195]), folate (SLC19 and SLC46 [196–198]), biotin and pantothenate (SLC5 [199]), nicotinate [200], thiamine (SLC19 and SLCO [201–204]), and riboflavin (SLC52 [205–209]).

In other cases the role of human protein transporters of natural products is well established, but their molecular nature (i.e. identity) has not yet been determined, e.g. those for psychoactive alkaloids such as cocaine [210] and nicotine [211–215] and opioids [216; 217]. Of course many transporters are known in the producer plants themselves [162–164].

Following this logic, the prediction, as tested here, is that at least some successful marketed drugs should be much closer to these plant and microbial molecules in structural terms than are the intermediary metabolites that are part of Recon2. **The prediction was amply demonstrated**, and serves to account for the otherwise anomalous finding that only a rather small fraction of intermediary <u>human</u> metabolites are reliably ‘similar’ (using the MACCS/TS metric) to marketed drugs at the level of 0.75 or 0.8. However, by contrast, we find that as many as 80% of natural products show such a similarity when surveyed extensively. This is consistent with the earlier, pilot findings (Fig 5C of [22]), and with the fact that Caco-2 permeability was poorly correlated with (the MACCS encoding of) endogenite-likeness [15].

### Evolutionary aspects of ‘secondary metabolites’ and other natural products

The question of what roles might be played by secondary metabolites in evolutionary terms is an old one, and almost certainly does not have a unitary answer. Note that the original definition of ‘secondary metabolites’ was to the effect that only a small number of organisms made a given such molecule [218]. However, most of this literature on secondary metabolites, taken as virtually synonymous with ‘natural products’, focuses on the benefits to be gained by the producer organisms <u>themselves</u>. To this end, there is abundant evidence that at least some natural products are used as signals by (and towards) other individuals of the same species, and are thus pheromones [219]. Necessarily, evolutionarily early variants of natural products may lack potency at the concentrations expressed [220] and the fact that the wider the number produced, the greater the likelihood of their selection [221] can explain why the selection pressure, <u>in terms of benefitting the producer,</u> may often be quite modest. However, our focus here is on the <u>benefits to consumer organisms</u>.

It now seems clear that our earlier focus [22–24; 114] on transporters just of human-encoded intermediary metabolites as the potential source of the ‘natural’ substrates of the transporters on which pharmaceutical drugs hitchhike was somewhat misplaced. This is because humans, and at least their vertebrate (and indeed invertebrate) evolutionary predecessors, have been exposed for millions of years to plant- and microbe-based dietary substances that had bioactivities of various kinds, many of which must have been beneficial and thus conferred a selective advantage, however small [222; 223], on the host. The origins of this remain uncertain, but the early ones (ca 1.8Gy BP) may have involved simple engulfment [224], with major eukaryote diversity set in place by 800My BP [225]. Even if one considers only angiosperms as potential sources of nutrients, these begin to arise ca 133 My BP [226] (with Solanaceae at around half that period BP [227]). Thus, the need (and ability) to take up plant and microbial metabolites is likely a trait of rather ancient origin.

### Transporter phylogenetics

Transporter phylogenetics is an area that is still highly under-researched (as are transporters in general [12]), and indeed nearly 100 new families are introduced into the Transporter Classification Database (TCDB) every year [228]. This is not the place to pursue that issue in detail, so a single example will suffice: a BLAST search of the sequence for human SLC22A4, the ergothioneine transporter, reveals (data not shown) that it is widespread among modern mammals, but obvious homologues are not to be found in reptiles, fishes, or lower taxa.

### Drugs and natural products

It is, of course, well known that many natural products can serve as medicines [87; 229], and that many purified substances derived therefrom are the basis of a significant fraction - probably 35-60% depending on how one counts - of marketed drugs (see above, and [157; 230–232]). We think that the present work highlights even more clearly how important natural products and their derivatives are likely to be in terms of producing novel, safe and efficacious drugs.

## Conclusions

The present analysis takes forward our continuing analysis of the structural similarities between marketed drugs and naturally occurring substances in two major ways. First, by looking at <u>rank orders</u> of similarities between encodings, we find very major differences, such that the metabolite with the closest TS to a drug in one encoding may be very different in both nature and TS value from that when compared with another encoding. There is no encoding that seems to us to have any special intellectual privileges, and as stressed by Everitt [233] unsupervised analyses should anyway best be judged simply on their utility. On this basis, we consider it entirely legitimate to pick and choose encodings to maximise apparent similarities, as a guide to testing, for instance, which other substances are competing substrates for the transport of a particular drug.

Previously, we focussed solely on human endogenites and the contents of Recon2 when making these comparisons. However, not least because of the discovery by GrUndemann and colleagues [64; 65] that SLC22A4 is in fact an ergothioneine transporter, we now recognise that we should include all kinds of plant- and microbial (and any other) natural products to which humans might have been exposed in evolution, and transport of whose bioactive principles might have been selected adaptively on the basis of their nutritional or medicinal activities and benefits. Clearly, natural evolution may be expected to have selected for the ability to transport molecules that in kind and amount were of benefit to the host. When we include such natural products, we find that the closeness of at least one of them, using one or more encodings, to marketed drugs is increased massively. This at once hereby points us at substances that might be the ‘natural’ substrates of a given drug transporter, suggests molecules for QSAR studies thereon, and potentially provides novel scaffolds for pharmaceutical drug discovery.

## Acknowledgements

We thank the BBSRC for financial support (grants BB/K019783/1 and BB/M017702/1), and Professor David Wishart and colleagues for providing their serum metabolome database in a particularly convenient format.

